# Pulmonary toxicity and inflammatory response of e-cigarettes containing medium-chain triglyceride oil and vitamin E acetate: Implications in the pathogenesis of EVALI but independent of SARS-COV-2 COVID-19 related proteins

**DOI:** 10.1101/2020.06.14.151381

**Authors:** Thivanka Muthumalage, Joseph H. Lucas, Qixin Wang, Thomas Lamb, Matthew D. McGraw, Irfan Rahman

## Abstract

Recently, there has been an outbreak associated with the use of e-cigarette or vaping products, associated lung injury (EVALI). The primary components of vaping products, vitamin E acetate (VEA) and medium-chain triglycerides (MCT) may be responsible for acute lung toxicity. Currently, little information is available on the physiological and biological effects of exposure to these products. We hypothesized that these e-cig cartridges and their constituents (VEA and MCT) induce pulmonary toxicity, mediated by oxidative damage and inflammatory responses, leading to acute lung injury. We studied the potential mechanisms of cartridge aerosol induced inflammatory response by evaluating the generation of reactive oxygen species by MCT, VEA, and cartridges, and their effects on the inflammatory state of pulmonary epithelium and immune cells both in vitro and in vivo. Cells exposed to these aerosols generated reactive oxygen species, caused cytotoxicity, induced epithelial barrier dysfunction, and elicited an inflammatory response. Using a murine model, the parameters of acute toxicity to aerosol inhalation were assessed. Infiltration of neutrophils and lymphocytes was accompanied by significant increases in IL-6, eotaxin, and G-CSF in the bronchoalveolar lavage fluid (BALF). In mouse plasma, eicosanoid inflammatory mediators, leukotrienes, were significantly increased. Plasma from e-cig users also showed increased levels of hydroxyeicosatetraenoic acid (HETEs) and various eicosanoids. Exposure to e-cig cartridge aerosols showed the most significant effects and toxicity compared to MCT and VEA. In addition, we determined at SARS-COV-2 related proteins and found no impact associated with aerosol exposures from these tested cartridges. Overall, this study demonstrates acute exposure to specific e-cig cartridges induces in vitro cytotoxicity, barrier dysfunction, and inflammation and in vivo mouse exposure induces acute inflammation with elevated pro-inflammatory markers in the pathogenesis of EVALI.

## 1. Introduction

Electronic Nicotine Delivery Systems (ENDS) products are battery-operated devices equipped with a tank, cartridge, or a pod filled with a liquid (e-liquid). These e-liquids may contain a base liquid (humectant), such as propylene glycol (PG) and vegetable glycerin (VG). Other contents include nicotine, flavoring chemicals, and flavor enhancers. Recently, cartridges containing tetrahydrocannabinol (THC) and cannabidiol (CBD) have been introduced to the ENDS market. These cartridges predominantly use carriers such as mineral oil and medium-chain triglycerides (MCT) oil. The heating element in these cartridges, i.e., coil/atomizer, raises the temperature of the e-liquid or oil to aerosolize its constituents. Some of these products have become abundantly available in the market containing adulterants, such as butane hash oil (a.k.a dabs) [1]. Using these products has caused acute lung injury to many e-cigarette (e-cig) users and the disease was recognized as e-cigarette, or vaping, associated lung injury (EVALI) in August 2019. As of February 2020, a total of 2807 hospitalized cases or deaths have associated with EVALI, according to the Centers for Disease Control and Prevention [2]. Symptoms of EVALI include labored breathing, dyspnea, chest pain, cough, nausea, diarrhea, fatigue, fever, and weight loss [3,4].

All EVALI patients have a history of e-cigarette use or vaping and the majority report using THC containing vaping products. No single constituent has been identified as the causative agent of EVALI. However, vitamin E acetate has been found in bronchoalveolar lavage fluid (BALF) of EVALI patients [5]. Most of the subsequent analyses have been geared towards the constituents of e-liquids like Vitamin-E acetate (VEA) as implicated in these ENDS-user subjects [6–8]. There is an urgent need for experimental-based studies to implicate the causative role of VEA in EVALI or other lung conditions reported. There is a gap in research and knowledge of pulmonary effects and vaping mediated toxicity. Additionally, inhalation toxicology studies in vitro and in vivo are needed to investigate the effects of exposure to constituents of e-liquids and vaping cartridges.

In this study, we provide the mechanisms of toxicity of inhalation exposure to e-cig cartridges and their major components. We assessed the total volatile organic compounds in the aerosols and conducted an in vitro toxicity assessment using lung epithelial cells and monocytes. The parameters of in vitro toxicity assessment included cell viability, ROS generation, trans-epithelial barrier function, and inflammatory mediators. To assess the acute inhalation toxicity of MCT, VEA, and e-cig cartridge aerosols in vivo, an acute exposure mouse model was utilized. In these exposed mouse groups, immune cell influx and cytokines in BALF were quantified to investigate the elicited inflammatory response compared to the unexposed counterparts. In addition, we studied at SARS-COV-2 related proteins and found no impact associated with aerosol exposures from these tested cartridges. We demonstrated a reduction in surfactant protein A in lung homogenates of VEA exposed mice, which plays a role in lipid homeostasis and innate immune defense. Lipidomics analyses performed on BALF of mice exposed to MCT, VEA, and e-cig cartridge aerosols, as well as plasma of e-cig users, showed changes in eicosanoids and, exhibiting their role in pulmonary inflammation. Further, we found altered glycerolipids, cholesterol esters, and glycerophospholipids in mouse BALF upon acute VEA and cartridge exposure. We hypothesized that VEA, MCT, and other chemical constituents present in e-cigarette cartridges [9] cause cellular toxicity, and oxidative and inflammatory responses in epithelial cells, monocytes, and in vivo in mouse lung upon inhalation.

## 2. Material and Methods

### 2.1 Scientific rigor and reproducibility

We have applied robust unbiased experimental design and data analysis approaches throughout the study. We have validated the methods and ensured reproducibility with repeated experiments. All methods are presented in detail with transparency. Results were reported and interpreted without bias. Laboratory grade biological and chemical resources were purchased from commercial sources. Our methodologies, data, and results adhered to strict NIH standards of reproducibility and scientific rigor.

### 2.2 Ethics statement: Institutional biosafety and animal protocol approval

Experiments in this study were performed according to the standards and guidelines approved by The University of Rochester Institutional Biosafety Committee (Study approval #Rahman/102054/09-167/07-186; identification code: 07-186; date of approval: 01/05/2019 and 02/03/2020). Validated cell lines, human bronchial epithelial cell lines (16-HBE and BEAS2B) and human monocytic leukemia derived cell line (MONO-MAC-6), were procured from ATCC, USA. Ethical approval was not necessary for the utilized cell lines.

All mouse housing, handling, exposure, and procedure protocols used in this study were approved by the University Committee on Animal Research (UCAR) Committee of the University of Rochester, Rochester, NY (UCAR protocol 102204/UCAR-2007-070E, date of approval: 01/05/2019 and 02/03/2020).

Human plasma samples used for lipidomics analysis were from the study conducted at the University of Rochester Medical Center (Rochester, NY, USA) (Institutional Review Board approval RSRB00064337) [10].

### 2.3 Aerosol exposure setup

An Ooze slim twist vaping pen set at 3.8V was connected to the Scireq Inexpose pump (Montreal, Canada). For in vitro exposures, cell culture plates were placed inside an Enzyscreen chamber (Enzyscreen BV, Netherlands) and exposed to two 70 mL puffs of the aerosol under air-liquid interface conditions. The cells were allowed to incubate in the vapor for 10-minutes.

For in vivo exposures, wild type mice with C57BL/6 background were exposed to 1 hr MCT, VEA, and cartridge aerosols with 70 mL puffs, two puffs/min using the Scireq Inexpose system (Montreal, Canada) [11].

### 2.4 Vitamin E acetate (VEA) 50% w/v preparation

A refillable vape oil cartridge was filled with either MCT oil (GreenIVe, Amazon) or tocopherol acetate (Sigma Cat# PHR1030-500MG) was mixed with MCT oil to make a 50% w/v or 1.06 M VEA.

### 2.5 E-cigarette cartridges

For the cell and mouse exposures in this study, e-cig cartridges used in our previous study characterizing constituents of EVALI were used [9].

### 2.6 Physicochemical characteristics of MCT and VEA

Respirable particle concentration and distribution:

Dusttrack II aerosol monitor 8530 (TSI, MN) was used to measure respirable particles with aerodynamic diameter 1.0, 2.5, 4.0, and 10.0 μm by taking readings immediately after a single puff was released to a chamber with dimensions 8”x6”x5.25” (Enzyscreen, Netherlands). Data collection duration is one minute with a 10s moving average. These fine particle measurements were obtained for air, MCT, VEA, and e-cig cartridge aerosols.

### 2.7 Measurement of total volatile organic compounds (VOC) levels in aerosols

To measure the VOC levels inside the Enzyscreen chamber, the exhaust tubing was connected to a 50 mL conical tube. A photoionization detector probe 985 (TSI, MN) was placed inside the conical tube, and the total VOC levels in air, MCT, VEA, and cartridge aerosols were recorded.

### 2.8 Acellular ROS Assay

Cell-free ROS generated by MCT, VEA, and e-cig cartridge aerosols were determined using 2’7’dichlorofluorescien diacetate (H2-DCF-DA) fluorogenic probe (Sigma-Aldrich Cat# 287810). H2-DCF-DA [5 mM] with NaOH [0.01N] was reacted for 30 minutes to prepare the dye. After incubation, 25 mM phosphate buffer was added to stop the reaction followed by the addition of 2 mL HRP. A set of hydrogen peroxide standards ranging from 0 to 50 μM was prepared from a 1M H2O2 stock. Subsequently, two puffs of MCT, VEA, and e-cig cartridges were bubbled through an impinger containing 10 mL of the dye connected to Scireq Inexpose pump. After aerosolization, the samples were incubated for 15 minutes at 37 °C in a water bath. Subsequently, the absorbance/emission was read at 485/535 nm with Turner Quantech fluorometer (FM109535, Barnstead international).

### 2.9 Cellular ROS Assay

2’7’dichlorofluorescien diacetate (H2-DCF-DA) fluorogenic probe (Sigma-Aldrich Cat# 287810) dye was prepared as described. 15,000 BEAS-2B cells/well were cultured in a 96-well plate in complete medium to 80% confluency. After serum depriving the cells at 1% FBS for 12 hours, 20 μM dye was added to each well of the 96-well plate and was incubated for 30 minutes. After incubation, the cells were treated with 0.25% (v/v) MCT, and 0.025% (w/v) or 50 μM VEA, and a mixture of e-cig cartridge liquids at 0.0001% (v/v). The absorbance/emission was read at 485/535 nm at 1.5 hours, 4 hours, and 6 hours using microplate spectrophotometer (Cytation 5, Biotek).

### 2.10 Cell Culture

Bronchial epithelial cells, BEAS-2B, were cultured in DMEM F-12 50/50 base media and supplemented with 5% FBS, 1% Pen/Strep, and HEPES. Cells were plated at 300,00 cells/well in 6 well plates in complete medium. At 80% confluency, the cells were serum-deprived in 1% FBS. Similarly, epithelial cells, 16-HBE, were cultured in DMEM base media and supplemented with 1% Pen/Strep, 10% FBS, and 1ml Amphotericin B. 16-HBE cells were plated and serum-deprived in 1% FBS. Monocytes, Mono-Mac-6 (MM6), were cultured in 24-well culture plates in RPMI 1640 media and supplemented with 1% Pen/Strep, 1mM Sodium Pyruvate, 2mM L-glutamine, non-essential amino acid, Transferrin, Oxaloacetic acid, Polymixin B, and Bovine Insulin. At 80% confluency, MM6 cells were serum-deprived in 1% FBS 12-hours before treatment.

### 2.11 Aerosol exposures and treatments to cells

Epithelial cells (BEAS-2B and 16-HBE) were exposed to MCT, VEA (50% w/v), and e-cig cartridges, as described in the exposure setup section. Twenty-four hours post-exposure, the conditioned media, and the cell pellets were collected for further analysis.

Monocytes, MM6 cells were treated with %0.25 (v/v) MCT, 0.025% (w/v) or 50 μM, and 0.0001% cartridges in 1% FBS media. Twenty-four hours later, conditioned media and cell pellets were collected.

### 2.12 Cytotoxicity Assay

To assess the induced cytotoxicity by MCT and VEA treatments, 20 μL of cells were mixed with 2OpL of Acridine Orange/Propidium Iodide. Then, 20 μl of the mixture was added to the Nexcelom automated cell counter slide, and viability and the live and total cell counts were performed using a Nexcelom 2000 Cellometer (Nexcelom Bioscience, Lawrence MA).

### 2.13 Cytokine ELISA

Conditioned media collected from each well was used to determine inflammatory cytokine release. Interleukin 6 (IL-6) was measured using an IL-6 ELISA kit following manufacturer’s instruction (Invitrogen, Catalog # CHC1263). Similarly, an interleukin-8 (IL-8) ELISA kit (Invitrogen, Catalog # CHC1303) to quantify secreted IL-8 levels in conditioned media. The cartridge aerosol exposed data points were then pooled, and the average was compared with air, MCT, and VEA.

### 2.14 Trans-epithelial electrical resistance (TEER) measurement

16-HBE cells (20,000 cells/well) were cultured on 24-well transwell inserts (Corning cat# 3470) until the cells reached a complete monolayer. Cells were then serum-deprived at 1% FBS 12-hours before exposure. Cells were exposed to MCT, VEA, and cartridge aerosols using the previously described aerosol exposure setup. 24-hours later, the barrier function was evaluated by recording the transepithelial voltage and resistance by EVOM2 (WPI instruments, FL). Each well was measured three times, and the average unit area resistance was calculated. Cartridge aerosol exposed data points were then pooled, and the average was compared with the unexposed air group.

### 2.15 In vivo mouse exposures

Approximately four month old male and female mice with C57BL/6 background were exposed to medium-chain triglyceride oil (MCT), vitamin E acetate (50% w/v in MCT), and e-cig cartridge aerosols one-hour per day for three consecutive days, as described in the aerosol exposure setup section [11]. For cartridge aerosols, six different cartridges were aerosolized for 10-minute cycles. Immediately following the last exposure, mice were euthanized and the tissues were collected.

### 2.16 Mouse arterial oxygen saturation

Immediately prior to mouse euthanasia, arterial oxygen saturation was measured in mice by MouseOX plus device (STARR life science, PA). Data collected over approximately ~8 minutes and any unstable data points were removed, including the first 60s.

### 2.16 Bronchoalveolar lavage (BALF) collection

Upon anesthesia 0.6 mL of 0.9% NaCl saline solution was instilled three times (1.8 mLs cumulative volume) into the trachea and the recovered BALF centrifuged at 1000 RPM for 7 minutes. The acellular fraction of the BALF was stored at −80°C for cytokine analysis by Luminex assay. The pelleted cells were then used for flow cytometry analysis to obtain differential cell counts.

### 2.17 Luminex assay

To determine inflammatory mediators released due to MCT, VEA, and e-cig cartridge aerosol exposures, 50 μL of the BALF sample was used with BioRad 23-plex-Group I kit (BioRad Cat# M60009RDPD) according to the manufacturer’s instructions. Briefly, capture antibody coupled magnetic beads were added to the plate, followed by the addition of samples and the standards. After incubating, detection antibody and streptavidin-PE were added. The appropriate number of washing steps and incubation steps were followed as instructed. After resuspending the sample in 125 μL of assay buffer, the plate was read on a FLEXMAP 3D system (Luminex). The concentrations of each analyte were compared to the unexposed air group and the analytes that showed significant differences were reported.

### 2.18 Flow cytometry analysis

Collected cells from the BALF recovery were counted by AO/PI assay to obtain total cell counts. The cells were then blocked with anti-CD16/32 (Fc block) for 10 minutes. Followed by a PBS wash step, cells were stained with CD45, F4/80, Ly6B.2, CD4, and CD8 cell surface markers in staining buffer to identify approximate counts of cell populations. After 30 minutes of incubation in the dark at 4°C the cells were washed twice in PBS and resuspended in 100 μL buffer. Appropriate FMO controls and compensation beads were used for compensation. Sample acquisition was performed using Guava easycyte 8 flow cytometer (Luminex). Data analysis was performed using GuavaSoft 3.3.

### 2.19 Western blot analysis

Snap-frozen mouse lung tissues were homogenized in RIPA buffer with protease inhibitor cocktail and 25μg of protein from air, VEA, MCT, and cartridge aerosol exposed mice (N=6 per group for male (n=3) and female (n=3) were extracted and quantified using BCA protein assay. Extracted proteins were separated on 7.5% SDS-PAGE gels, which were then transferred onto nitrocellulose membranes. The membrane was probed with SP-A antibody (ab115791), ACE2 antibody (ab108252), Furin (ab183495), and TMPRSS2 (ab92323). B-actin (ab20272) was used as a loading control for normalization.

### 2.20 Lipidomics analysis

Mouse BALF samples (200 μL) and human plasma from one of our e-cig studies were analyzed by Cayman chemicals, MI, for eicosanoids/oxylipins and short-chain-fatty-acids by LC-MS/MS. Heat maps were generated depicting the changes in analytes. Analytes showing distinct differences were then graphed as box-whisker plots. Untargeted lipidomic profiling was performed by normalizing peak areas with internal standards. Air control samples were compared with VEA and cartridge exposed samples, and only the statistically significant analytes were included in **Table 1.**

**Table 1:** Untargeted lipidomic analysis (DG: Diradylglycerols, CE: Sterols, PC: Glycerophosphocholines) in bronchoalveolar lavage fluid from air, vitamin E acetate (VEA), and cartridge aerosol exposed mice. Normalized peak area means ± SEM with P-value and significance vs. air using two-way ANOVA (*p<0.05, **0.01<p and ***p<0.001 vs. Air, P>0.05 are denoted as NS, N=4/group).

| Analyte | Air<br>Mean | ±SEM | VEA<br>Mean | ±SEM | P-value | Significance<br>(vs. Air) | Cartridge<br>Mean | ±SEM | P-value | Significance<br>(vs. Air) |
| --- | --- | --- | --- | --- | --- | --- | --- | --- | --- | --- |
| DG(16:0_18:2_0:0) | 06.49 | ±0.20 | 05.05 | ±0.36 | <0.001 | *** | 04.38 | ±0.04 | <0.001 | *** |
| DG(16:0_16:0_0:0) | 00.19 | ±0.01 | 00.25 | ±0.04 | 0.5741 | NS | 00.40 | ±0.04 | 0.0031 | ** |
| CE(20:4) | 02.77 | ±0.10 | 02.52 | ±0.09 | 0.0002 | *** | 02.87 | ±0.24 | 0.2751 | NS |
| PC(16:0/16:0) | 29.86 | ±2.18 | 31.48 | ±2.41 | <0.001 | *** | 40.82 | ±2.64 | <0.001 | *** |
| PC(16:0_16:1) | 16.77 | ±1.42 | 17.59 | ±0.85 | 0.0295 | * | 23.46 | ±1.17 | <0.001 | *** |
| PC(16:0_18:2) | 04.31 | ±0.28 | 05.02 | ±0.48 | 0.0644 | NS | 06.74 | ±0.75 | <0.001 | *** |
| PC(16:0_18:1) | 04.63 | ±0.44 | 04.88 | ±0.17 | 0.6911 | NS | 06.64 | ±0.41 | <0.001 | *** |
| PC(14:0_16:0) | 04.22 | ±0.37 | 04.59 | ±0.34 | 0.4375 | NS | 05.79 | ±0.58 | <0.001 | *** |
| PC(16:0_22:6) | 01.86 | ±0.18 | 02.33 | ±0.32 | 0.2937 | NS | 03.18 | ±0.51 | <0.001 | *** |
| PG(16:0_18:1) | 02.70 | ±0.27 | 02.93 | ±0.11 | 0.7163 | NS | 03.85 | ±0.21 | 0.0014 | ** |
| PG(18:2_16:0) | 02.36 | ±0.16 | 02.84 | ±0.22 | 0.2688 | NS | 03.93 | ±0.37 | <0.001 | *** |
| PG(16:0/16:0) | 01.73 | ±0.14 | 01.85 | ±0.07 | 0.9170 | NS | 02.58 | ±0.12 | 0.0237 | * |

### 2.21 Oil-Red-O staining

Oil-Red-O staining was performed on lung OCT-cryosections and immobilized MM6 cells using BioVision Cat# K580-24 according to manufacturer’s instructions. In brief, the oil-red-o stock was made in 20 mL 100% isopropanol. The slides were incubated in 60% isopropanol for 5 minutes, followed by evenly covering with Oil-Red-O working solution. The slides were then placed on a shaker and incubated for 20 minutes. After rinsing the slides with dH2O, hematoxylin was added and incubated for 1 minute. After rinsing the slides with dH2O five times, the slides were viewed under the microscope. Lipid-laden-indices were calculated in representative images of MM6 stained with Oil-Red-O.

### 2.22 Lipid-laden-index (LLI) scoring

The method for LLI scoring of the Oil-Red-O stained monocytes was adapted from Kazachkov et al. [12]. In two representative images, the total macrophages in each field were counted. Then, macrophages with <50% of the cytoplasm opacified by the stain were assigned a score of “1”. Macrophages with >50% of the cytoplasm opacified by the stain were assigned a scored of “2”. The LLI was then calculated as follows: LLI=((% 1+ macrophages)×1)+((% 2+ macrophages)×2).

### 2.23 Statistical Analysis

Statistical analysis of data was done by One-Way ANOVA with Tukey’s multiple comparison test for multiple sample groups with one variable and by Two-Way ANOVA with Tukey’s multiple comparison test for multiple sample groups with two variables using GraphPad Prism 8.0. Data are reported by mean ± SEM and statistical significance was reported as *p< 0.05, **p<0.01, and ***p<0.001.

## 3. Results

### 3.1 MCT, VEA, and e-cig cartridges induce cellular and acellular ROS generation and cytotoxic responses

BEAS-2B cells generated increased cellular ROS production upon treatment with MCT, VEA, and cartridges compared to untreated controls. Cellular ROS levels increased with respect to time. The highest cellular ROS generation was observed in MCT-treated cells, followed by VEA-treated and cartridge-treated cells (N=4) (Figure 1A).

**Figure 1:**
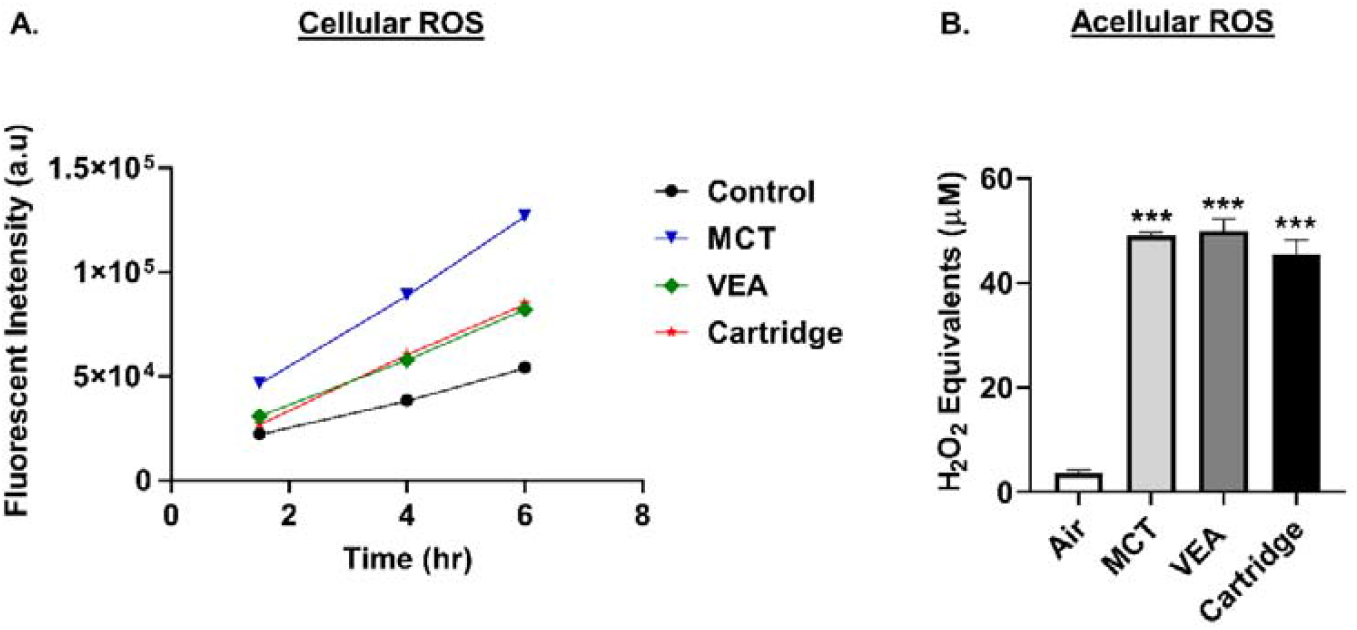
Increased cellular and aceullar generation by MCT, VEA, and e-cig cartridges. (A) Increased cellular ROS generation by cartridge liquid treatments. BEAS-2B cells were seeded at 15,000/well. The following day, cells were treated with 20 μM DCFH-DA in 1% FBS containing media for 30 min. Cells were then washed and treated with 0.25% (v/v) MCT, 0.025% (w/v) or 50 μM Vitamin E acetate, and e-cig cartridge liquids. Fluorescence (485,535nm) was read at 1.5,4 and 6h post-treatment (N=4). Control vs. MCT ***p<0.0001. Control vs. VEA *p<0.05, Control vs. Cartridge p=0.2236. (B) Increased aceullar generation by MCT, VEA, and e-cig cartridges. Air, 50% (w/v) vitamin E acetate, MCT, and e-cig cartridges were bubbled through dichlorofluorescein diacetate (DCFH-DA) dye with two 70 mL puffs. Fluorescence was measured at 485/535 nm. One-way ANOVA ***p<0.001 vs. Air (N=2-4).

Acellular ROS was determined by drawing vapors from VEA, MCT, and e-cig cartridges through DCFH dye. MCT, VEA, and e-cig cartridges generated significantly increased acellular ROS levels (in H2O2 equivalents) compared to air, but did not differ between treatment groups One-way ANOVA ***p<0.001 vs. Air (N=2-4) (Figure 1B).

BEAS-2B cells exposed to VEA and e-cig cartridge aerosols caused variable cell death, although non-statistically significant, compared to the air group. Among aerosols, e-cig cartridges caused the greatest cytotoxicity (up to 34% cell death, n=6). Monocytes, MM6, treated with VEA or MCT did not cause significantly increased cytotoxicity. However, MM6 treated with 0.25% e-cig cartridge liquid caused significant cytotoxicity (up to 98%, n=9).

### 3.2 MCT, mineral oil, VEA, and e-cig cartridge aerosols contain volatile organic compounds

The highest total VOCs were released by the vape cartridges (20.03 ppm) followed by MCT (10.33 ppm), then mineral oil (7.33 ppm), and VEA (9.67 ppm) compared to air (0.00 ppm).

### 3.3 Exposure to MCT, VEA, and e-cig cartridges elicited a differential inflammatory response in epithelial and monocyte cells

Conditioned media from aerosol exposed cells were assayed for IL-8 and IL-6 to determine the elicited inflammatory response by individual cells. Both MCT and VEA induced a mild increase in IL-8 and IL-6 compared to their unexposed controls. Exposure to e-cig cartridge aerosols induced highly significant IL-6 and IL-8 levels (One-way ANOVA, *p<0.05 and ***p<0.001 vs. control, n=6) (Figure 2A,B).

**Figure 2:**
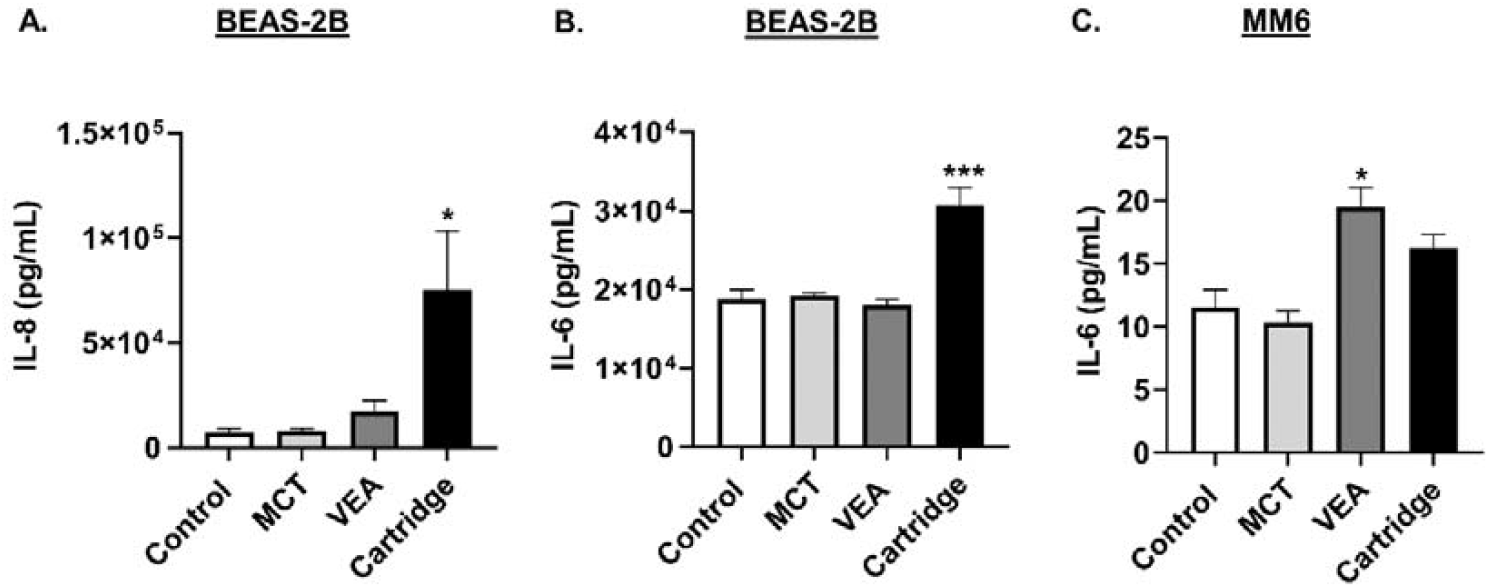
Induction of differential cytotoxicity and inflammatory cytokines, IL-8 and IL-6, responses by e-cig cartridge exposure in lung epithelial cells and monocytes. BEAS-2B cells exposed to MCT (medium-chain triglycerides) oil, vitamin E acetate (50% w/v in MCT), and e-cig cartridges with 2 puffs of 70 mL in total 10 minutes. 24 hours later, cytokines (A) IL-8 and (B) IL-6 were measured in the conditioned media by ELISA. One-Way ANOVA *p<0.05 and ***p<0.001 vs. control. MCT and VEA are nonsignificant vs. control (N=6/group). (C) Mono-Mac-6 cells were seeded at a density of 5×105 cells/well in 5% FBS containing media. The following day, cells were treated with 50 μM vitamin E acetate, 0.25% MCT, and e-cig cartridges at 0.25%. 24-hours post-treatment IL-6 cytokine level was measured in the conditioned media by ELISA. One-Way ANOVA *p<0.05, **p<0.01, and ***p<0.001 vs. control. P>0.05 were considered nonsignificant and were not denoted with asterisks (N=3-9).

In MM6 cells, while the treatment with MCT did not cause any changes in these cytokine levels, treatment with VEA and cartridge liquid caused an elevation in IL-6 levels. VEA induced significantly elevated IL-6 levels in monocytes (One-way ANOVA, *p<0.05 vs. control, n=6) (Figure 2C).

### 3.4 Reduced epithelial barrier function following exposure to MCT, VEA, or e-cig cartridges

Trans-epithelial electrical resistance (TEER) decreased significantly in 16-HBE with the treatment of all three aerosols: MCT, VEA, or e-cig cartridge. The unexposed monolayer of epithelial cells’ TEER was approximately 68 Ω.cm2 (68.13± 6.39; n=3) while MCT, VEA, and e-cig cartridge exposure resulted in a significant decrease in the TEER measurement to 40.2 Ω.cm2,40.1 Ω.cm2, and 38.8 Ω.cm2, respectively (One-way ANOVA, p***<0.01 vs. air, N=3) (Figure 3).

**Figure 3:**
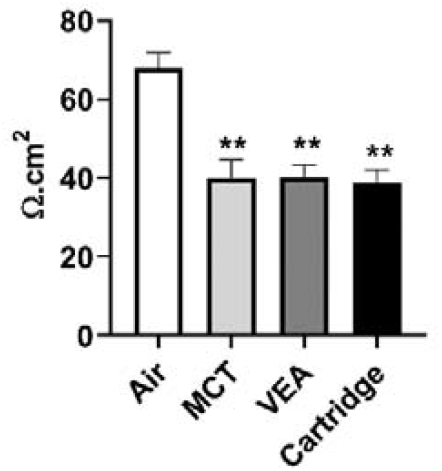
Barrier dysfunction caused by e-cig cartridges in bronchial epithelial cells. 16-HBE cells were cultured on transwell inserts and exposed to 2 puffs of 70 mL of air, MCT, vitamin E acetate (50% w/v), and e-cig cartridge for 10 minutes. 24-hours later, the resistance was measured by EVOM2 device. One-Way ANOVA **p<0.01 vs. Air (N=3).

### 3.5 Treatment of macrophages with MCT, VEA, and e-cig cartridge liquids caused varying levels of lipid-laden macrophage formation

Treatment with MCT, VEA, or e-cig cartridges showed various levels of lipid-laden macrophage formation. In representative images, treatment with MCT, VEA, and cartridges resulted in lipid-laden indices of 56.7±32.9, 1.2±1.6, and 30.8±3.7, respectively, compared to 3.4±4.8 for untreated controls, (N=2) (Figure 4).

**Figure 4:**
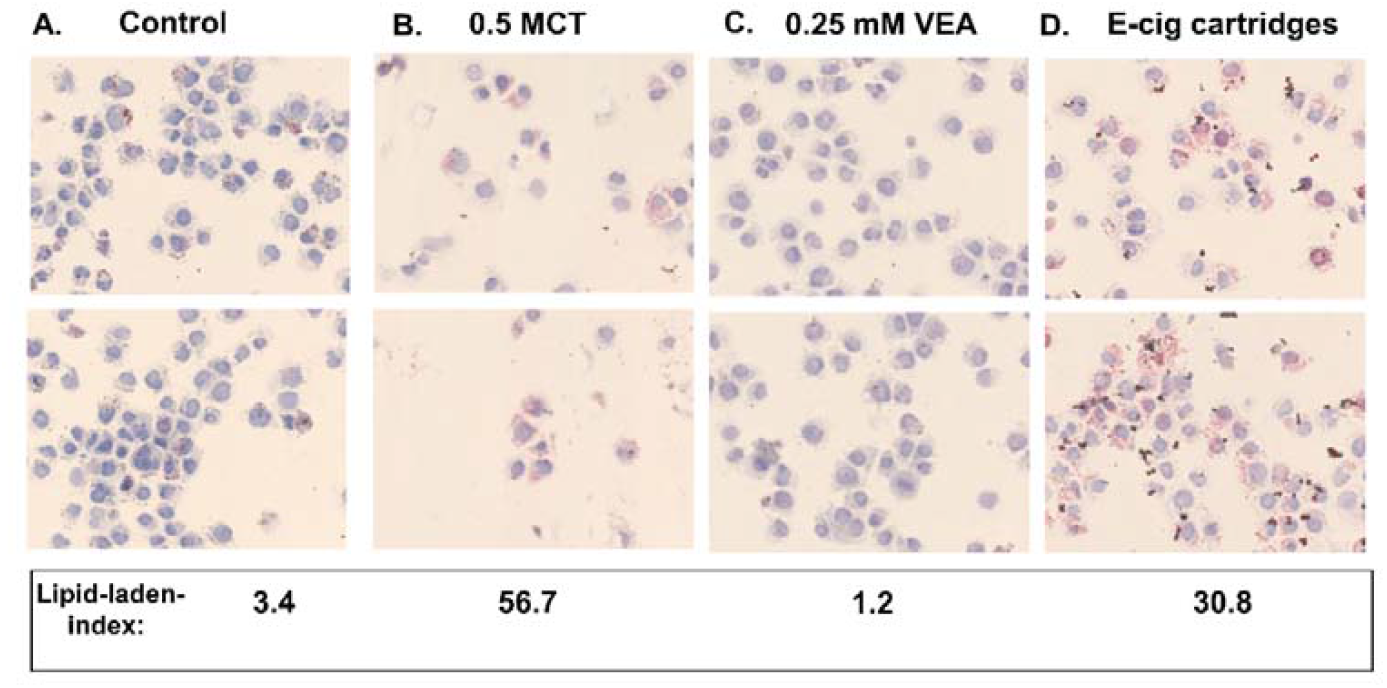
Representative images of lipid-laden macrophages formed by MCT, VEA, and e-cig cartridge liquid treatments. MM6 cells were seeded in 5% complete media and treated with 0.00005% (0.0003%) six unknown cartridge liquids 6h later. 24h post-treatment cells were collected and stained with Oil-Red-O (N=3). The averaged lipid-laden indices of the representative images reported.

### 3.6 Acute exposure to e-cig aerosols did not alter arterial oxygen saturation in mice

Mouse arterial oxygen saturation was approximately 96%, 92%, 94%, and 95% for air, MCT, VEA, and cartridge aerosol exposed mice, respectively.

### 3.7 Exposure to e-cig cartridge aerosols induced an inflammatory response in mice

Acute exposure to e-cig cartridge aerosols caused increased bronchoalveolar lavage fluid (BALF) total cells (one-way ANOVA *p<0.05 vs. air, N=6) (Figure 5A). Neutrophils and CD4-lymphocytes in the BALF increased by 5.1-fold and 6.9-fold, respectively, but these increases were not statistically significant (Figure 5B,C).

**Figure 5:**
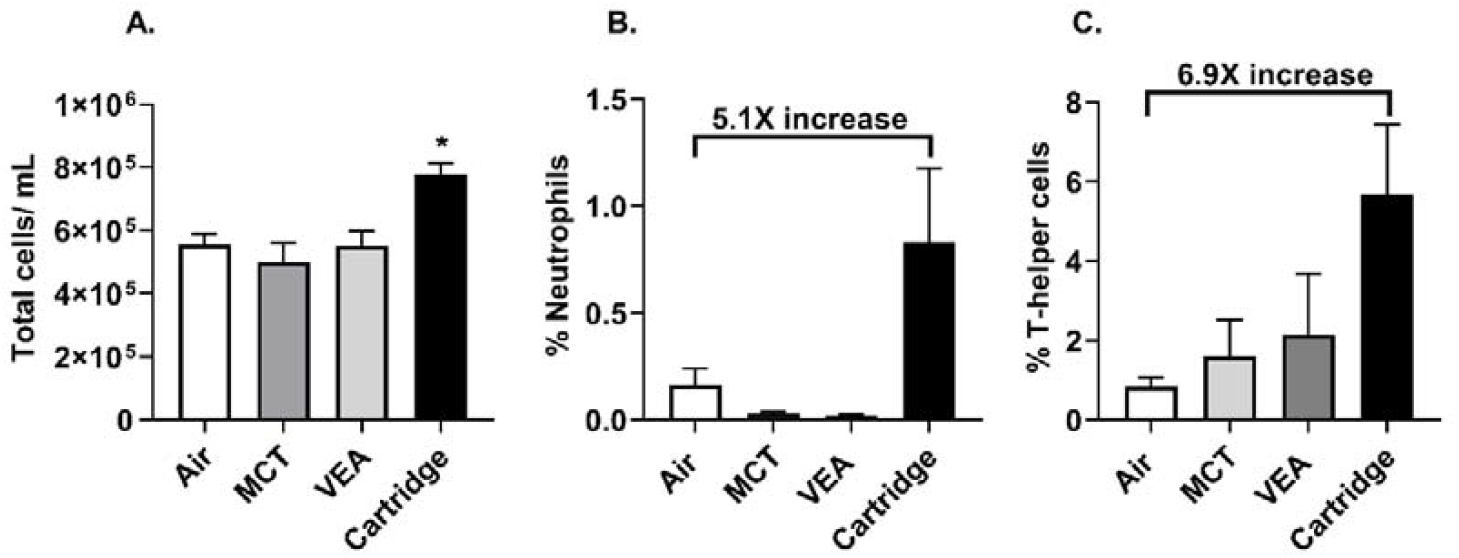
Exposure to aerosols from e-cig cartridges induced a differential immune cell influx. C57BL/6 background wild type mice were exposed to MCT, vitamin E acetate 50% w/v, and six unknown cartridges with one puff a minute, 1 hour/day for three consecutive days. Mice were sacrificed immediately after the last exposure, and the immune cells in the BALF were assessed. (A) Total cells in air, MCT, VEA, and e-cig cartridge exposed BALF. One-Way ANOVA *p<0.05 vs. Air (N=5). (B) Neutrophil count in BALF (N=5). (C) CD4+ cell count in BALF (N=5). P>0.05 vs. air were considered nonsignificant and were not denoted with asterisks.

Acute exposure to VEA and cartridges resulted in increased BALF inflammatory mediators, IL-6, and eotaxin (one-way ANOVA *p<0.05, ***p<0.001 vs. air, N=6) (Figure 6A,B). Granulocyte colony-stimulating factor (G-CSF) was also significantly increased in mice exposed to e-cig cartridge aerosols, but not with MCT or VEA aerosols (one-way ANOVA *p<0.05 vs. air, N=6) (Figure 6C). Exposure to MCT, VEA, and cartridges resulted in significantly attenuated levels of MCP-1, RANTES, IL-17A, IL-12p40, and IL-4 (one-way ANOVA *p<0.05, **p<0.01, and ***p<0.001 vs. air, N=6) (Figure 6D-H).

**Figure 6:**
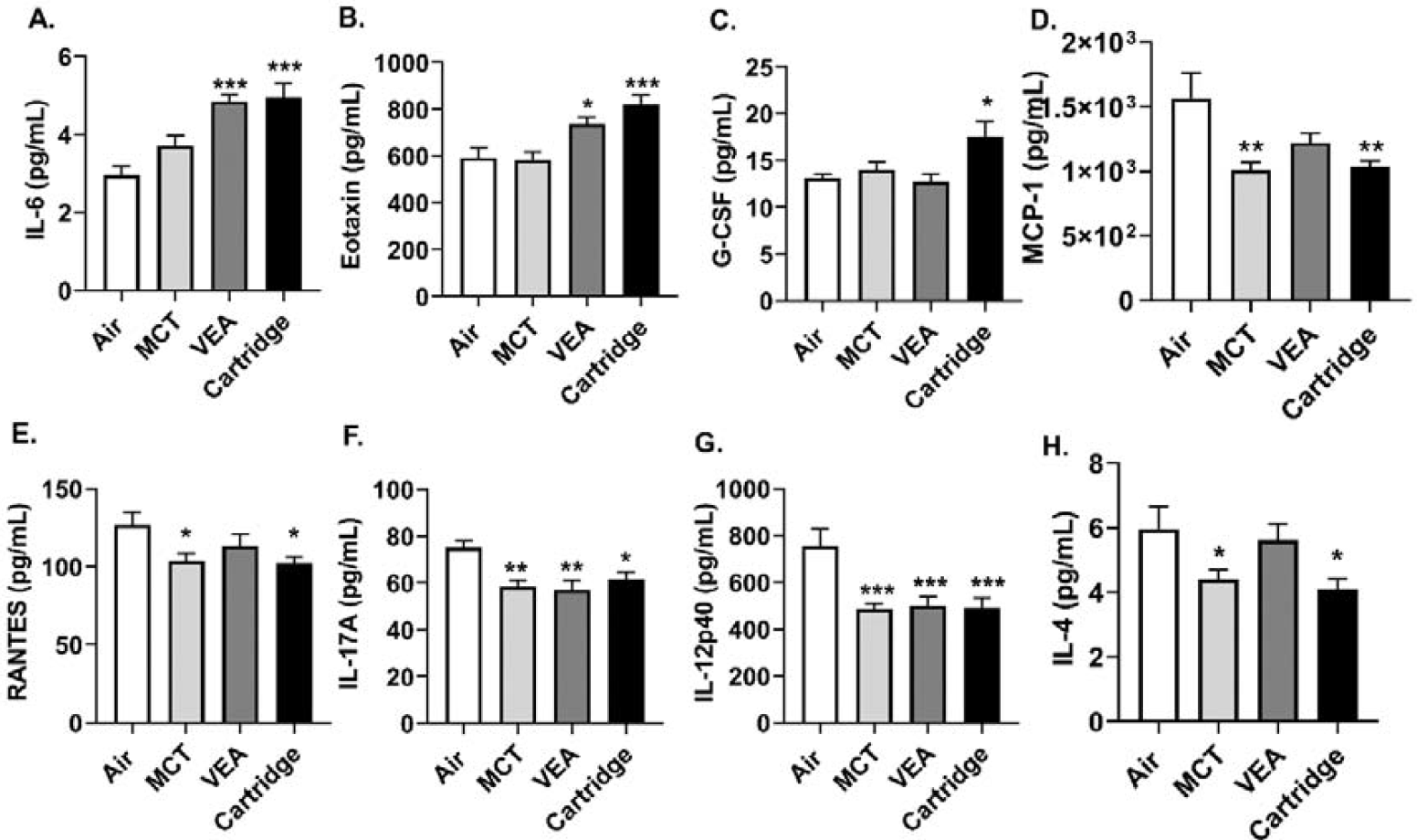
Induction of differential cytokine response by MCT, VEA, and e-cig cartridge aerosol exposure in mouse BALF. Wildtype mice with C57BL/6 background were exposed to MCT, vitamin E acetate 50% (w/v), and e-cig cartridges, 1 puff/min for 1 hour for three consecutive days using Scireq inExpose system. Immediately after the last exposure, the mice were sacrificed and the BALF was collected. Secreted inflammatory mediators were measured using BioRad 23-plex, group 1 Luminex kit. (A) IL-6, (B) Eotaxin, and (C) G-CSF were (D) MCP-1, (E) RANTES, (F) IL-17A, (G) IL-12p40, and (H) IL-4 are reported. One-Way ANOVA *p<0.05, **p<0.01, and ***p<0.001 vs. Air, P>0.05 were considered nonsignificant and were not denoted with asterisks (N=6).

### 3.8 Surfactant-associated protein A (SP-A) reduced in VEA exposed mouse lung homogenates

Lung homogenates from mice exposed to air, VEA, MCT, and e-cig cartridge aerosols were quantified for lung surfactant protein-A (SP-A). There was a significant reduction in SP-A in lung homogenates from male mice exposed to VEA. This difference was not seen in female mice (Figure 7A,B).

**Figure 7:**
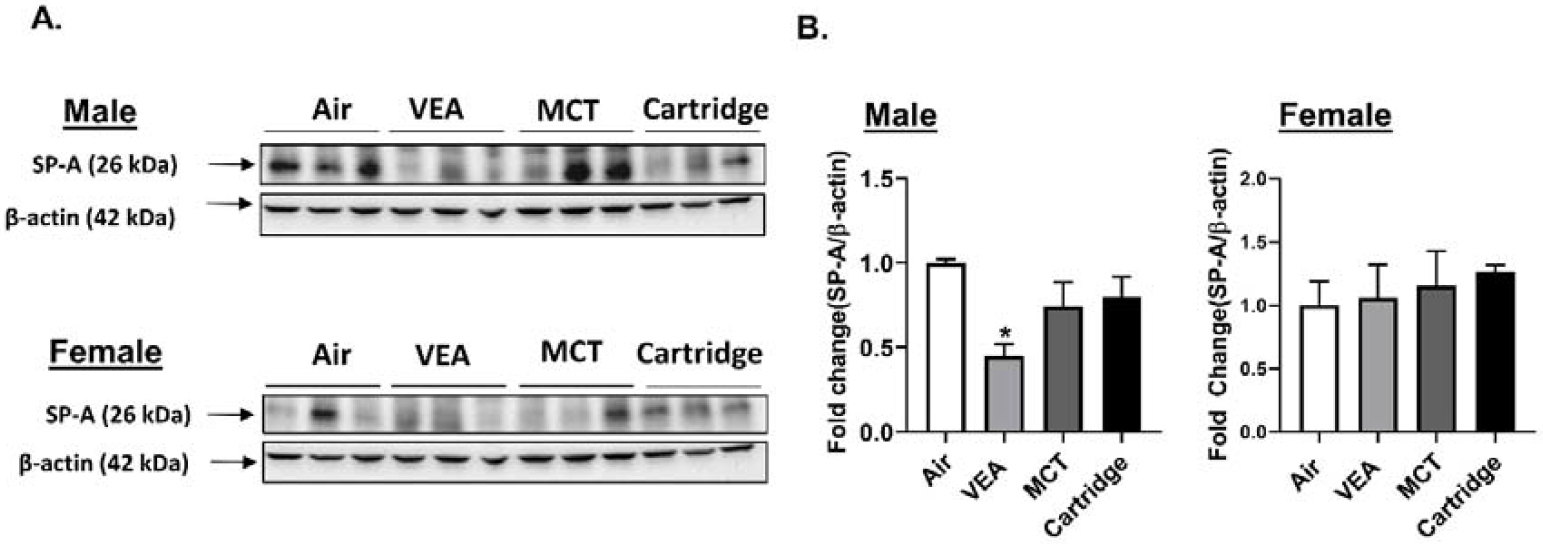
Surfactant-associated protein A (SP-A) was reduced in VEA exposed mouse lung homogenates. (A)Surfactant-associated protein A (SP-A) protein abundance in mouse lung homogenates from C57Bl/6 background male and female mice was determined by immunoblot analysis. (B) The quantified results are expressed as means ± SEM. Significance was determined by 1-way ANOVA with Bonferroni’s correction for multiple comparisons. **P < 0.05 vs. Air. P>0.05 were considered not significant and were not denoted with asterisks (n = 3 per group for either sex).

### 3.9 Differential changes in eicosanoids/oxylipins and short-chain fatty acids in mouse BALF following VEA and e-cig cartridge aerosols exposure

Exposure to VEA caused changes in eicosanoids, including 6kPGF1a, LTB4, LTD4, LTE4, and 5HETE. Exposure to e-cig cartridge aerosols increased LTB4, LTC4, LTD4, LTE4,5HETE, and 12HETE. LTC4 levels were significantly elevated with VEA exposure compared to the unexposed air group (one-way ANOVA ***p<0.001 vs. air, N=4) (Figure 8A,C). No significant changes in short-chain fatty acids were detected (Figure 8B).

**Figure 8:**
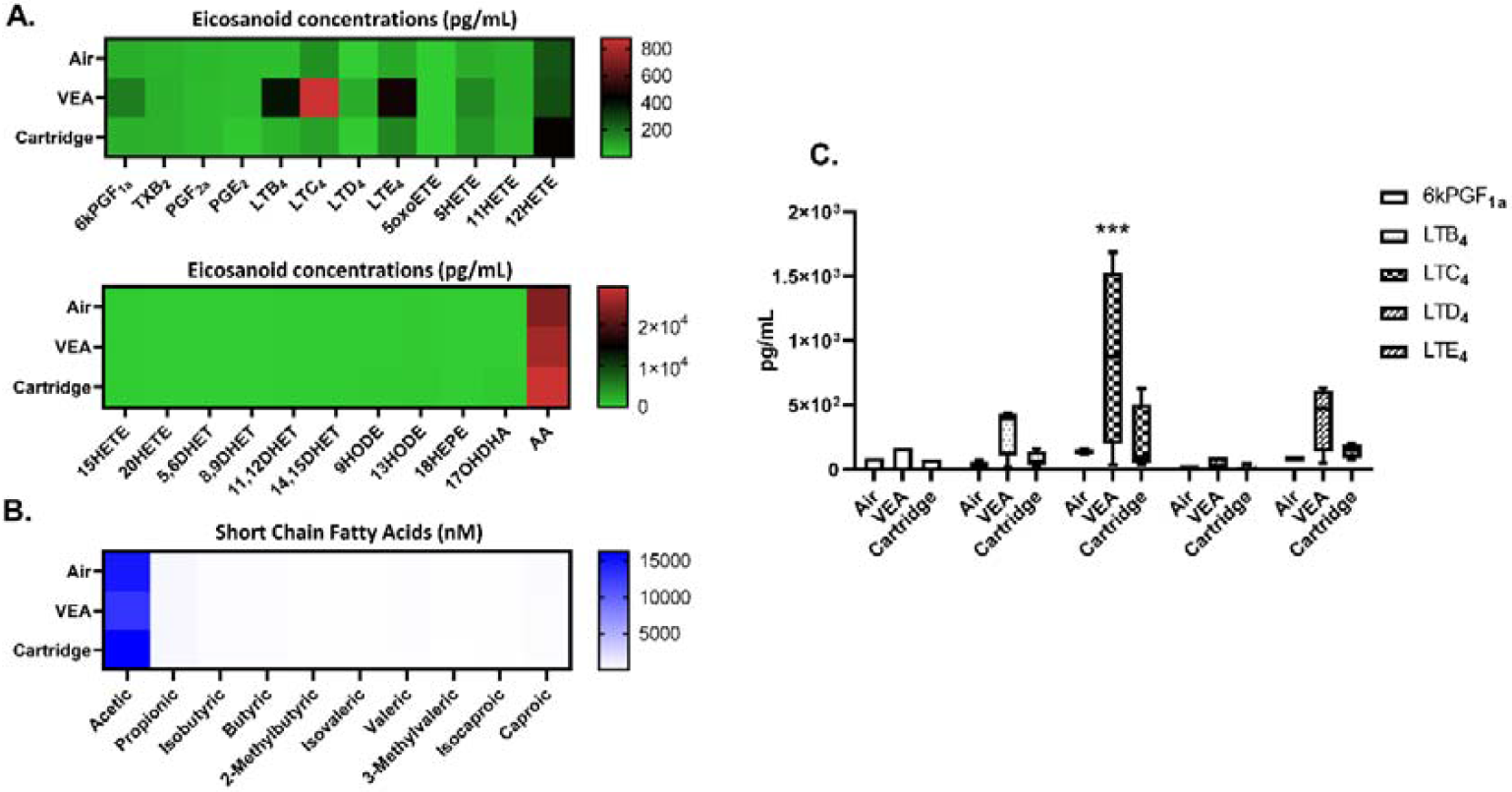
Eicosanoids, oxylipins, short-chain fatty acids in bronchoalveolar lavage fluid from VEA, and e-cig cartridge exposed mice. Heat map generated from relative quantification by LC-NS/MS of (A)eicosanoids, oxylipins, and (B) short-chain fatty acids in BALF from mice exposed to VEA and unknown cartridges. (C) analytes, 6KPGF1a, LTB4, LTC4, LTD4, and LTE4 in box-whisker plots ***p<0.001 cs. Air (N=4/ group). P>0.05 were considered nonsignificant and were not denoted with asterisks.

### 3.10 Diradylglycerols (DG), sterols (CE), and glycerophosphocholines (PC) were signifincatly altered in VEA and cartridge aerosol expose mice

Lipidomic profiling on aerosol exposed mouse BALF showed changes in glycerolipids (diradylglycerols), cholesterol esters (sterols), and glycerophospholipids (glycerophosphocholines and glycerophosphoglycerols). As shown in **table 1**, DG(16:0_18:2_0:0), CE(20:4), PC(16:0/16:0), and PC(16:0_16:1), levels were significantly altered with VEA exposure in BALF. With the cartridge aerosol exposure apart from the listed above DG(16:0_16:0_0:0), PC(16:0_18:2), PC(16:0_18:1), PC(14:0_16:0), PC(16:0_22:6), PG(16:0_18:1),PG(18:2_16:0), and PG(16:0/16:0) levels were significantly altered in mouse BALF.

### 3.11 E-cig users exhibited differential changes in eicosanoids/oxylipins and short-chain fatty acids in human plasma

E-cig users had less TXB2 and AA and increased levels of 5HETE, 11HERE, 12HETE, and 17OHDHA levels in their plasma compared to nonsmokers (N=6) (Figure 9 A, C, D). Reduced acetic add levels were also observed in e-cig users compared to nonsmokers (N=6) (Figure 9B).

**Figure 9:**
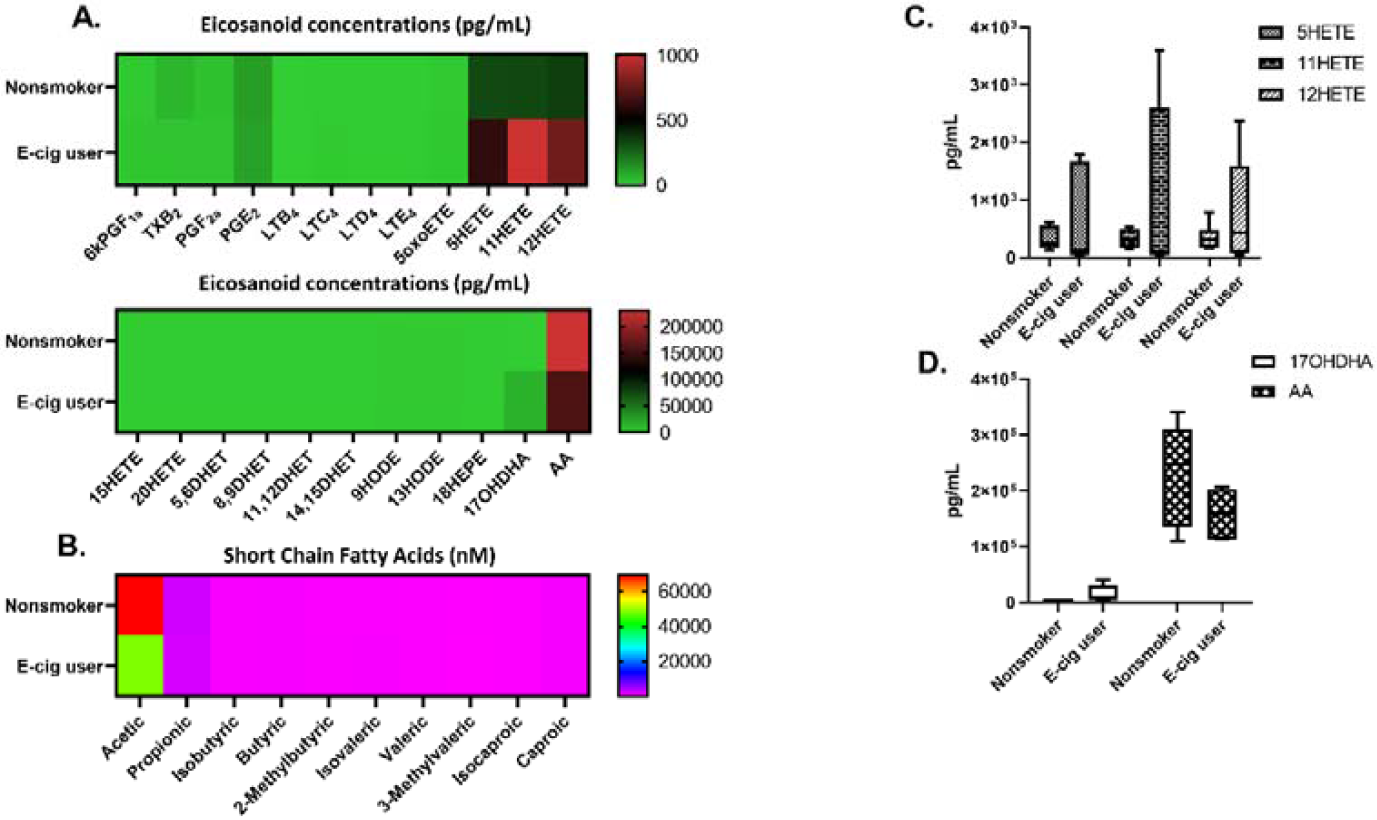
Plasma eicosanoids, oxylipins, short-chain fatty acids in nonsmokers and e-cig users. (A) Heat map generated from relative quantification by LC-NS/MS of eicosanoids, oxylipins, and (B) short-chain fatty acids in plasma from nonsmokers and e-cig users. (C) 5HETE, 11HETE, and 12HETE levels in plasma (D) 17OHDHA and AA in plasma (N=6/ group). P>0.05 vs. nonsmoker were considered insignificant and were not denoted with asterisks.

### 3.12 SARS-COV-2 proteins ACE2, TMPRSS2, and Furin were largely unaffected by cartridge aerosols

Both male and female mice exposed to VEA and e-cig cartridge aerosols showed no difference in ACE2, TMPRSS2, and furin. Male mice exposed to MCT showed a significant decrease in spike protein cleaving protease, furin (Figure 10 A,B).

**Figure 10:**
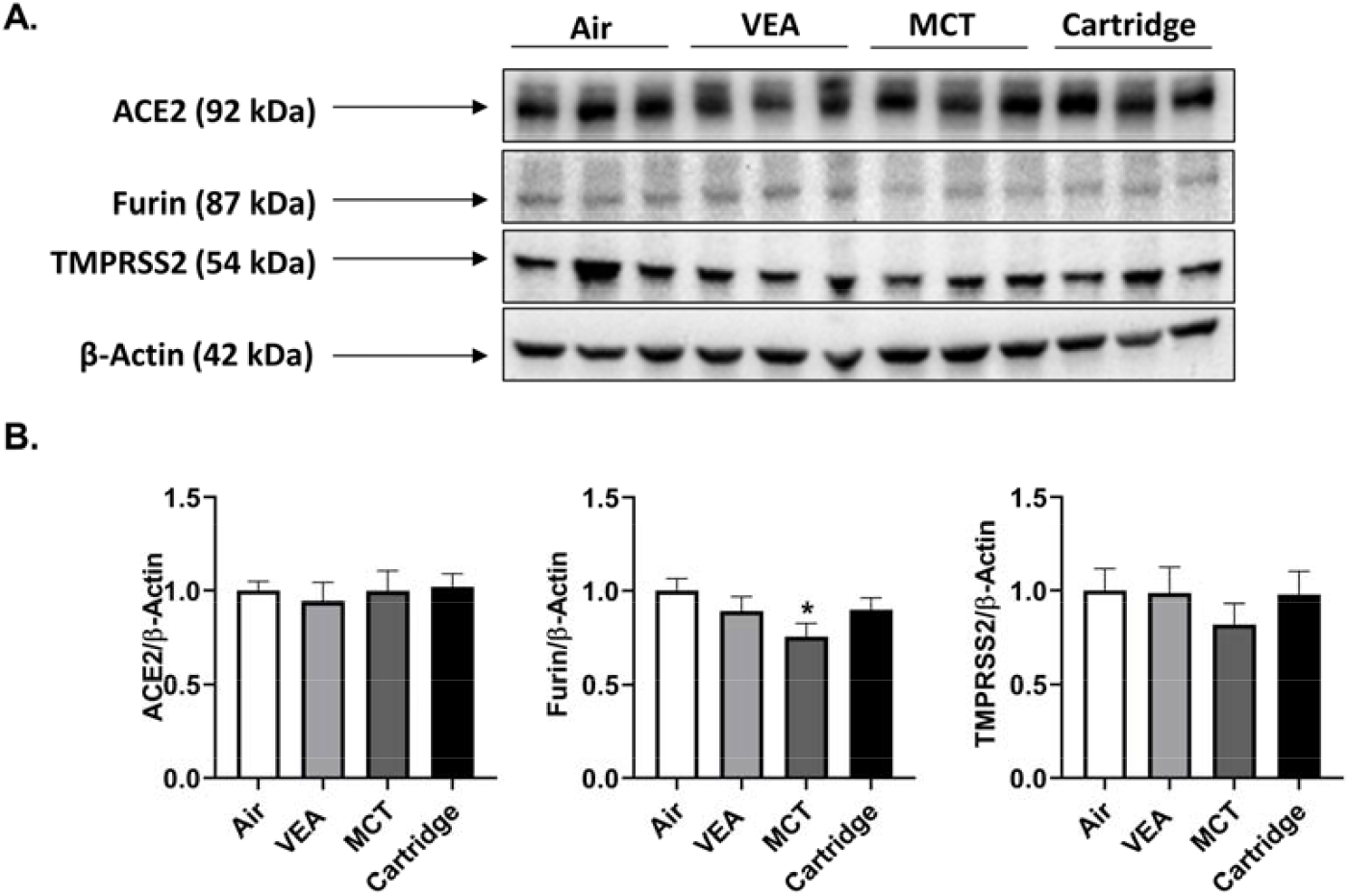
SARS-COV-2 spike protein cleaving enzyme, Furin, was reduced in MCT exposed in lung homogenates of male mice. After the three-day acute exposure to MCT, VEA, and cartridge aerosols, the abundance of ACE2, Furin, and TMPRSS2 proteins were determined in mouse lung homogenates by immunoblotting assay. (A) Immunoblots for Ace2, Furin, TMPRSS2, and β-actin. (B) Data represented as fold-change ±SEM *p<0.05 vs. air. P>0.05 were considered insignificant and were not denoted with asterisks (One-way ANOVA, N=3/group for either sex).

### 4. Discussion

Exposure to certain electronic nicotine delivery systems (ENDS) cartridges has recently increased hospitalizations due to respiratory failure [6–8,13,14]. According to case reports since 2012, e-cig users with symptoms without direct etiology have undergone various diagnoses, including acute lung injury, atypical pneumonitis, and eosinophilic or lipoid pneumonia [6,13]. The presence of abnormal lipid-laden macrophages in lung tissue samples from EVALI subjects has been associated with lipoid pneumonia [15,16]. The patients diagnosed with lipoid-pneumonia suffered from cough, difficulty in breathing, shortness of breath, chest tightness and pain, nausea, vomiting, fatigue, fever, and weight loss. In one of our studies, we analyzed the chemical constituents of e-cig cartridges involved in EVALI [9]. Users of these ENDS products developed varying degrees of respiratory illnesses, but the mechanism of toxicity of developing EVALI is still unknown.

In this study, we studied the mechanisms of toxicity in developing EVALI. Constituents in e-cig cartridges affect the inflammatory state of pulmonary epithelial cells, immune cells, and exposed mice. MCT and mineral oil are used as the vehicle (humectant) in these cartridges. Upon heating, these products emit volatile organic compounds. In cartridge aerosols, we saw nearly double the amount of VOCs compared to MCT, mineral oil, or VEA aerosols. The highest total VOCs were released by the vape cartridges followed by MCT oil, mineral oil, and VEA compared to air, suggesting carbonyl stress by these compounds. These VOCs in e-cig cartridges can generate toxic compounds, such as THC/VEA carbonyl complexes and ketene, during pyrolysis, leading to lung damage [17,18]. In this study, we also observed significant acellular and cellular ROS generation by cartridges and their major components. These highly reactive species vary based on vaping patterns and can lead to lung pathogenesis [19]. Increased levels of IL-6 and IL-8 are clear indicators of an ongoing inflammatory response in epithelial cells and monocytes. IL-8 is a potent neutrophil attractant that can induce lung damage by degranulation of stored mediators and enzymes, oxidative bursts, as well as the release of neutrophil extracellular traps [20,21].

Exposure to e-cig cartridge aerosols showed severe toxicity in monocytes, as well as significantly elevated IL-6 levels, a biomarker for lung injury. This indicates that in less than 24 hours, the immune system can be primed towards a pro-inflammatory response [22]. In this study, two puffs of aerosols significantly affected the epithelial barrier function; this is consistent with other studies our lab, as well as other researchers, have published [23,24]. Toxicants emitted in these aerosols damage tight junctions between the epithelial cells disrupting the epithelial barrier. This alteration of the epithelial permeability drives pathogenesis by promoting inflammatory signaling pathways [23,24].

Radiological and histological analyses of lungs with darkened patches and ground-glass opacities suggest that volatile hydrocarbons such as terpenes and oils in e-cigs are possibly involved in the pathogenesis of EVALI [25,26]. Various hydrocarbons and reactive aldehydes are formed upon heating and thermal decomposition of e-liquids around 500 °F. All these cartridges are used at around a 3.5V to 5.5 V setting using a specific device, which is similar to CBD containing cartridges. These emitted chemicals may play an important role in the pathogenesis of EVALI, apart from MCT and VEA. Further studies with chronic animal exposures are required for histopathological studies.

Formation of foamy macrophages in BALF and lung tissue has been used as a diagnostic tool for vaping associated lung injuries [16,26,27]. In this study, we observed the formation of lipid-laden macrophages in MM6 cells treated with MCT, VEA, and cartridges. E-cig cartridge liquids showed more Oil-Red-O staining compared to VEA, suggesting that contents of these cartridges, such as other oils, including MCT, can potentially cause exogenous lipoid pneumonia upon inhalation and aspiration [28]. In our short 1 hr x 3 day exposure, we noticed some Oil-Red-O staining in BALF macrophages; however, we did not notice any lipid-laden macrophages in lung sections (data not shown), suggesting that the formation of these lipids may take longer than this exposure duration. In the cytokine profile, significantly increased IL-6, eotaxin, and G-CSF were observed. Increased G-CSF in BALF has been seen in patients with ARDS (acute respiratory distress syndrome), which has been correlated with pulmonary neutrophilia [29–31]. This is consistent with our findings that showed neutrophil influx (Figure 5A). In many cases, acute alveolitis has been associated with an influx of eosinophils and neutrophils [32,33]. Similar to radiological images of lungs showing ground-glass opacity, presence of lipids, and the influx of cells usually seen in cases with hydrocarbon and oil inhalation have also been identified in images from vaping related lung injuries [34]. The increase in eotaxin (Figure 6B) suggests eosinophilic chemotaxis in the lung tissue and lavage as eotaxin plays a major role in recruiting eosinophils to the sites of inflammation [35]. Corroborating our data, exposure to ENDS has been shown to augment the levels of IL-6 in BALF along with increased neutrophils and CD4+ cells [14].

In many studies, IL-6 has been shown to increase with exposure to any insult causing acute lung injury to maintain the homeostasis of the inflammatory response [36–38]. The increase in G-CSF and eotaxin along with the increase in CD4 cells in BALF suggest that acute exposure may be leaning toward the Th2 pathway, implicating potential eosinophilic pneumonia with prolonged exposure [39–41]. Excessive inflammation results in lung damage in diseases. IL-17 plays an important role in leukocyte infiltration. Studies have shown reduced G-CSF secretion in IL-17R-/- mice and that these mice were protected against ARDS [42,43]. In this study, aerosol exposure resulted in attenuated levels of MCP-1, RANTES, IL-17A, IL12p40, and IL-4 in BALF (Figure 6D-H). The occurrence of eosinophilia and eosinophil alveolitis has been observed in patients with idiopathic pulmonary fibrosis [44]. Consistent with observations in this study, increased CD3+ lymphocytes in BALF and protection against fibrosis in IL-4 deficient mice have been observed in another study [45]. Moreover, RANTES and MCP-1 are heavily involved in leukocyte influx and bronchial hyperresponsiveness [46]. Similarly, the observed significant decrease in IL12p40 suggests the balancing of the propagated inflammatory response as IL12p40 can be pro-inflammatory and pro-fibrotic [47]. The reduction of these cytokines and chemokines implicates a negative feedback of the immune system to sustain homeostasis. In a previous study, we observed that cannabidiol (CBD) containing e-cigarettes possess anti-inflammatory properties by upregulating the MCPIP1 transcription factor [48]. As the tested e-cig cartridges possibly contained THC and CBD, it is possible that these drugs also contributed to the induced cytokine storm. Further studies are needed to understand the mechanisms of this immune modulation, such as understanding the role of innate lymphoid cells (ILCs) in response to exposure to e-cigarette cartridge components and toxicants. ILC2 and ILC3 are pivotal to the homeostatic response to environmental toxicants with a poised immune response to alleviate lung damage and maintain lung function [49].

In our lipidomics analysis of mouse BALF exposed to VEA and cartridge aerosols, we observed increased levels of lipid mediators: leukotrienes. Leukotrienes (LTCs) are 5-lipoxygenase metabolites that are vital for lung epithelial and endothelial barrier function [50]. Cysteinal leukotrienes, such as LTC4, have been shown to play a role in allergic asthma development via group 2 innate lymphoid cells (ILC2) activation and propagation [51]. In other animals such as dogs, rats, and rabbits, increased LTC4 in BALF has been seen in association with injured lungs and the formation of extravascular lung water [52]. These animal studies suggest that in our murine model, exposure to VEA and cartridges has promoted allergic asthma-related inflammation-causing lung injury, especially in VEA exposed mice. However, it may be possible that these lower LTC4 levels seen in the cartridge exposed group may be confounded by the presence of nicotine as nicotine has LTC4 suppressive effects [53,54],

In e-cig users of our cohort, we observed increased systemic oxidative stress and inflammatory responses [10]. These samples also showed increased hydroxyeicosatetraenoic acids (HETEs) in plasma. HETES are well-known biomarkers of oxidative stress, and elevated levels of HETE isomers have been found in smokers [55]. In this study, we found HETE isomers 5, 11, and 12 induction in e-cig users’ plasma with other lipid mediators. These metabolites of arachidonic acid play an important role in mediating inflammation in the lungs. HETES are involved in neutrophil transcellular migration and proliferation of cells elicited inflammatory response during acute lung injury [56,57]. Moreover, our untargeted lipidomics analysis showed altered glycerolipids (DG), cholesterol esters (CE), and glycerophospholipids (PG and PC) in BALF. A study comparing metabolomic similarities between smokers and cigarette smoke-exposed mice revealed highly correlated augmented levels of glycerolipids and glycerophospholipids in mouse BALF with the exposure to cigarette smoke [58]. Thus, it can be inferred that a similar modulation in lipids occurred with cartridge exposures. Surfactant-associated protein A (SP-A) was reduced in lung homogenates of VEA exposed male mice. Surfactant proteins are important to prevent lung atelectasis and the reduction of SP-A may suggest excess accumulation of lipids in the alveolar space as we demonstrated with this acute exposure. Consistent with our observations, reduced surfactant proteins in BALF and lung homogenates have been seen in mice exposed to ENDS independent of nicotine [14].

Amid the COVID-19 outbreak, smokers have shown to have more adverse outcomes upon infection. In our acutely exposed lung homogenates, we looked at ACE2, TMPRSS2, and Furin protein abundance and observed no association with cartridge exposures. The decrease in furin, spike glycoprotein cleaving protease, in male mice with the MCT exposure needs to be further investigated[59]. The exacerbated response in smokers and vapers may suggest that smoking/vaing with nicotine and α7nAChR with ACE2 may be involved [60].

Overall, based on our data, it is possible that both VEA and e-cig cartridge may have caused acute respiratory distress inducing eosinophilic pneumonia. As we found in our previous study, most of these cartridges contained THC derivatives and VEA [9]. At low concentrations, VEA was not cytotoxic. Moreover, VEA has been employed as a carrier in therapeutic interventions [61], suggesting that it may act as a carrier for THC in the blood and the brain. Upon aerosolization, VEA or its formed oxidant derivatives may interact with phospholipids and surfactants of the epithelial lining fluid, thus adversely affect the normal lung function [62,63]. In e-liquids, VEA is used along with mineral oil and MCT oils, and there’s a lack of understanding of inhalation toxicity of these additives and diluents. Based on our results, MCT had lesser effects compared to VEA in vivo, but it showed comparable effects to VEA in vitro; overall, cartridges containing both of these components showed the greatest effects. Other than these two primary components, it may be possible that other constituents in cartridges are also responsible for EVALI; those chemicals include cyclooctasiloxane, cyclohexasiloxane, cyclotrisiloxane, tricaprylate, decanoic acid, benzoic acid, 2,2-dimethoxybutane, and tetramethyl silicate [9].

In summary, this study demonstrates acute exposure to specific e-cig cartridges induce in vitro cytotoxicity, barrier dysfunction, and inflammation and in vivo mouse exposure induces acute inflammation with elevated pro-inflammatory markers. It was also found that SARS-COV-2 related proteins had no impact associated with aerosol exposures from these tested cartridges or other agents tested. Overall, this study suggests that prolonged exposure to these ENDS may cause significant lung damage, which is involved in the pathogenesis of EVALI.

## 5. Acknowledgments

This study was supported by NIH 1R01HL135613, WNY Center for Research on Flavored Tobacco Products (CRoFT) # U54CA228110, and toxicology training program grant T32-ES007026. We thank Cayman Chemicals for lipidomics analyses. We are thankful to Isaac K. Sundar, Ph.D. for his assistance in performing animal sacrifices and his scientific input. We thank Samantha McDonough, BS and Krishna Maremanda, Ph.D. for their technical support and scientific contribution. We thank Gary Ginsberg, Ph.D. at the New York State Department of Health, Center for Environmental Health, Albany, NY, for assisting the study design, providing scientific input throughout this study, and for editing the manuscript.

## 6. Author Contributions

TM and IR: Conceived and designed the experiments. Wrote and edited the manuscript.

TM, TL, JL, and QW: Assisted in writing the manuscript, conducted experiments, performed data analyses.

MDM: Provided scientific input and edited the manuscript.

## 8. Data availability statement

We declare that we have provided all the data, but the primary data will be available upon request.

## 9. Disclaimer

The authors have nothing to claim or disclaim about any products used here to test their toxicological and biological effects. The authors have no personal interests or gains from the outcome of this study. The products tested are available commercially to consumers/users.

